# Antibody-mediated depletion of human CLEC-2 in a novel humanised mouse model

**DOI:** 10.1101/2021.10.03.462933

**Authors:** Helena C Brown, Sarah Beck, Stefano Navarro, Ying Di, Eva M Soriano Jerez, Jana Kaczmarzyk, Steven G. Thomas, Valbona Mirakaj, Steve P Watson, Bernhard Nieswandt, David Stegner

## Abstract

Platelet C-type lectin-like receptor 2 (CLEC-2) has been proposed as a potential anti-thrombotic target as genetic or antibody-mediated receptor deficiency prevents occlusive thrombus formation in mice. This occurs through interaction with an unknown ligand as the endogenous ligand podoplanin is not present in the vasculature. However, the CLEC-2-podoplanin interaction does have an important role in tumour metastasis. There are currently no methods to test potential human therapeutics targeting CLEC-2, such as antibodies, *in vivo*. We have therefore generated and characterised a humanised CLEC-2 mouse (hCLEC-2^KI^) and developed a novel monoclonal anti-human CLEC-2 antibody, HEL1, for *in vivo* testing. hCLEC-2^KI^ mice were phenotypically normal and had comparable platelet glycoprotein receptor expression, activation and aggregation to wildtype platelets. hCLEC-2^KI^ mice had both comparable bleeding and vessel occlusion times to WT mice. Challenging hCLEC-2^KI^ mice with HEL1 or a second monoclonal anti-hCLEC-2 antibody, AYP1, resulted in transient thrombocytopenia as well as CLEC-2 depletion for more than 2 weeks but had no effect on haemostasis. This illustrates the power of the humanised CLEC-2 mouse model in evaluating novel therapeutics *in vivo*, including antibodies that target CLEC-2, as well as the limited effect on haemostasis when targeting CLEC-2.

## Introduction

C-type lectin-like receptor 2 (CLEC-2) is a unique platelet activation receptor signalling through a single YXXL sequence, representing half of an immunoreceptor tyrosine-based activation motif (hemITAM).^1,2^ CLEC-2 and its endogenous ligand podoplanin are crucial for normal development with mice deficient in either showing defective blood-lymphatic vessel separation.^3-5^ This interaction also has a role in tumour metastasis.^6,7^

CLEC-2 is important in thrombosis, particularly in thrombus stability, and immuno-thrombosis^2,8-10^ with CLEC-2 deficiency reducing vessel occlusion in several *in vivo* thrombosis models with little effect on haemostasis.^11-13^ Occlusion is unaltered in CLEC-2 Y7A signalling null mice, in which the receptor is normally expressed, suggesting it is the presence of CLEC-2 itself, rather than CLEC-2-induced platelet activation, that has a role in thrombus stability.^12^ Furthermore, immunodepletion of CLEC-2 from the platelet surface using the monoclonal antibody INU1 has similar effects on thrombus formation, with depletion lasting up to 6 days and accompanied by transient thrombocytopenia.^8,11^

As a result, CLEC-2 has been suggested as a potential anti-thrombotic target. However, the *in vivo* role of human CLEC-2 cannot be readily investigated experimentally in humans, meaning there are limited methods to test potential therapeutics pre-clinically. In addition, although antibodies against human CLEC-2, such as AYP1,^14^ exist it is unknown if human, like mouse, CLEC-2 can be immunodepleted. Here we have generated a humanised CLEC-2 mouse model that can be used to test potential anti-human CLEC-2 therapeutics *in vivo*.

## Methods

### Animals

Humanised CLEC-2 (hCLEC-2^KI^) mice were generated by replacing the mouse *Clec1b* gene on chromosome 6 with the corresponding region of the human *Clec1b* gene using CRISPR/Cas9 (supplemental Figure 1). Mice were genotyped by PCR using the following primers: fwd: 5’-GCAAAACAAACCCCAAGTGTCCTGG-3’, WT-rev: 5’-ATGCCCAAATTGCTGAATGAGCCTT-3’ and KI-rev: 5’-CCGTTATCCCCTTGACTTCTATGCCC-3’ yielding a 247 bp PCR product for the WT allele and a 479 bp product for the KI allele. Animal experiments were approved by the district government of Lower Franconia.

### Blood donors

Blood samples were obtained from healthy volunteers after obtaining written informed consent in accordance with the Declaration of Helsinki and with approval from the Institutional Review Boards of the University of Wuerzburg.

### Antibody Generation

CLEC-2 was immunoprecipitated from human platelet lysates using Protein G Sepharose beads coupled to AYP1 and used to repeatedly immunise female Wistar rats. Splenic B-cells were fused with Ag14 myeloma cells and HAT medium was used to select for hybridomas. These were tested by flow cytometry for secretion of hCLEC-2 specific antibodies; supernatant from each hybridoma was incubated with hCLEC-2^KI^ mouse blood before washing in 1 ml PBS by centrifugation. The supernatant was discarded and the blood was incubated with anti-rat IgG-FITC before being analysed on a BD FACS Calibur. Positive hybridomas were subcloned and tested for hCLEC-2 specificity twice before monoclonal antibody purification.

### Antibodies and Reagents

The mouse anti-human CLEC-2 antibodies AYP1 and AYP2, and the rat-anti mouse CLEC-2 antibodies INU1 and INU2, as well as recombinant mouse podoplanin-FC, were generated in house as previously described^8,14^ whereas the AYP1-FITC conjugate was purchased from Biolegend. Anti-mouse and anti-rat IgG-FITC antibodies were from Dako and the GAPDH antibody from Sigma Aldrich. All other antibodies were from Emfret Analytics. ADP, apyrase, prostacyclin (PGI_2_), fibrinogen and haematoxylin were from Sigma Aldrich. HORM collagen was from Nycomed Pharma, thrombin from Roche and U46619 from Alexis Biochemicals; heparin was from Ratiopharm, eosin was from Roth. Protein G Sepharose was from GE Healthcare. Rhodocytin and collagen related peptide (CRP) were generated as described previously.^15,16^

### Mouse Platelet Preparation

Mice were bled from the retro-orbital plexus into heparin using a heparinised glass capillary. Blood was centrifuged twice at 300 *g* for 6 min to obtain the platelet-rich plasma (PRP) which was transferred into additional heparin prior to the second centrifugation. 0.02 µg/ml apyrase and 0.1 µg/ml prostacyclin were added to the PRP, which was further centrifuged at 800 *g* for 5 min to pellet the platelets. Platelets were resuspended in Tyrode’s buffer supplemented with apyrase and prostacyclin, counted and centrifuged again before being resuspended at 5 × 10^5^ for aggregation or 3 × 10^5^ platelet/µl for spreading assays and left to rest at 37°C for 30 min prior to use.

### Human Platelet Preparation

Citrated blood was taken from healthy volunteers and further anticoagulated with acid citrate dextrose (ACD). PRP was obtained by centrifugation at 300 *g* for 20 min and 0.02 µg/ml apyrase, 0.1 µg/ml prostacyclin and 100 µl/ml ACD were added before centrifugation at 500 *g* for 10 min. The resulting platelets were resuspended in 2 ml Tyrode’s buffer supplemented with apyrase, prostacyclin and 150 µl/ml ACD and further centrifuged at 500 *g* for 10 min. Platelets were counted and resuspended in Tyrode’s buffer at 5 × 10^5^ platelet/µl and left to rest at 37°C for 30 min prior to use.

### Platelet Aggregation

Washed platelets were further diluted with Tyrode’s buffer supplemented with 2 mM Ca^2+^ and 100 µg/ml human fibrinogen for all agonists except thrombin where Tyrode’s buffer without fibrinogen was used. Platelet aggregation was measured using a 4-channel aggregometer (APACT) under stirring conditions for 10 min after the addition of 5 µg/ml collagen, 0.01 U/ml thrombin, 0.24 µg/ml rhodocytin, 1-20 µg/ml HEL1 or 10 µg/ml of either INU1 or AYP1 antibodies or HEL Fab fragments.

For aggregation using ADP PRP was obtained as described above but without the addition of extra heparin. It was diluted in Tyrode’s buffer supplemented with 2 mM Ca^2+^ and aggregation was measured for 10 min after the addition of 5 µM ADP.

### Platelet Spreading

Platelets were allowed to spread at 37°C for 30 min on fibrinogen and CRP or 60 min on podoplanin before being fixed. For spreading on fibrinogen 0.01 U/ml thrombin were added to the platelets immediately prior to spreading. Platelets were imaged using a Zeiss Axiovert 200 microscope in DIC mode fitted with a 100x objective. 5 fields of view per coverslip were imaged and the platelets were categorised into 4 groups: 1) unspread platelets, 2) platelets forming filopodia, 3) platelets with lamellipodia and 4) fully spread platelets.

### Western Blots

Mouse and human platelets were prepared as described above and resuspended at 1 × 10^6^ platelets/µl in lysis buffer prior to centrifugation at 20,000 *g* for 10 min at 4°C to remove cell membranes. Lysates were mixed with reducing (AYP2) or non-reducing (AYP1, INU2 and HEL1) Laemmli buffer. Samples were separated by SDS-PAGE and transferred onto PVDF membranes which were blocked and then incubated with primary antibody. Membranes were incubated with secondary HRP-conjugated antibodies which were detected using ECL and an Amersham Imager.

### Platelet Count

Blood was collected in EDTA coated tubes and platelet count was measured using a ScilVet analyser.

### Flow Adhesion Assay

Coverslips were coated with 200 µg/ml HORM collagen overnight at 37°C then blocked in 1% BSA. Blood was diluted 1:2 in Tyrode’s buffer supplemented with 2 mM Ca^2+^ and platelets were labelled with a Dylight 488 conjugated anti-GPIX antibody. Blood was perfused over the coverslips at a shear rate of 1200 s^-1^ for 4 min and washed for a further 4 min with Tyrode’s buffer supplemented with calcium. 8 representative fields of view were imaged in both brightfield and fluorescence using a Leica DMI6000B microscope with a 63x objective. Images were then analysed in terms of platelet surface coverage and volume (fluorescent integrated density) using Fiji.^17^

### Flow Cytometry

#### Glycoprotein Expression

50 µl of mouse blood was collected in heparin, further diluted 1:2 in PBS and incubated with saturating amounts of anti-GPIb, GPVI, GPV, GPIbβ, GPIX, αIIbβ3, β3, α2, α5, mCLEC-2 or hCLEC-2 FITC-conjugated antibodies. 500 µl of PBS were added prior to measuring surface expression on a BD FACS Celesta. For assessing CLEC-2 expression on human platelets whole blood was diluted 1:20 in PBS and incubated with 2 µl AYP1-FITC.

#### Platelet Count

Blood was incubated with saturating amounts of anti-GPIX-FITC and anti-αIIbβ3-PE conjugated antibodies and double positive platelets were counted for 30 s.

#### Platelet Activation

Blood was diluted with 1 ml of Tyrode’s buffer and centrifuged twice at 800 *g* for 5 min, then resuspended in 750 µl of Tyrode’s supplemented with 2 mM Ca^2+^. Saturating amounts of JON/A-PE and anti-P-selectin-FITC antibodies were added as well as 10 µM ADP, 3 µM U46619, 0.001 – 0.1 U/ml thrombin, 0.1 – 10 µg/ml CRP, 1.2 µg/ml rhodocytin or 10 µg/ml AYP1 and incubated for 15 min. 500 µl PBS were added and platelet activation measured.

### CLEC-2 Depletion

3 µg/g bodyweight of either AYP1, HEL1 or INU1 were injected intraperitoneally and platelet count and CLEC-2 surface expression were measured for up to 25 days. Antibody binding to platelets was determined using anti-mouse (AYP1) or anti-rat (HEL1 and INU1) IgG-FITC antibodies; prior to incubation, blood was further diluted in 1 ml PBS and centrifuged at 800 *g* for 5 min to remove any unbound CLEC-2 antibody. Mice were depleted 5 to 10 days before *in vivo* experiments.

### Tail Bleeding Time

Mice were anaesthetised by intraperitoneal injection of medetomidine (0.5 μg/g; Pfizer, Karlsruhe, Germany), midazolam (5 μg/g; Roche, Grenzach-Wyhlen, Germany) and fentanyl (0.05 μg/g; Janssen – Cilag, Neuss, Germany). 1.5 mm of the tail was cut with a scalpel and blood was collected every 20 s using filter paper without touching the wound. This was done until bleeding ceased up to a maximum of 20 min.

### Mechanical injury of abdominal aorta

6-12 week old mice were anaesthetised as described above and the abdominal aorta was exposed. A Doppler ultrasonic flow probe (Transonic Systems) was placed around the aorta and thrombosis was induced upstream by mechanical injury resulting from a 15 s compression using forceps. Blood flow was monitored until either it ceased (complete vessel occlusion for 1 min) or 30 min had elapsed.

### Histology

Intestine, brain, liver, spleen, lymph nodes and femora were removed and fixed before being dehydrated and embedded in paraffin. Sections were stained with haematoxylin and eosin and imaged using a Leica DMI400B microscope. Megakaryocytes were counted in spleen and femoral sections across 8 fields of view.

### Statistical analysis

Normally distributed data, as determined using a Shapiro-Wilk test, are shown as mean ± standard deviation and were analysed by ANOVA followed by multiple comparison correction where appropriate, unless otherwise stated. Data that do not follow a normal distribution are shown as median with interquartile range and were analysed by Mann-Whitney tests unless otherwise stated.

## Results and Discussion

hCLEC-2^KI^ mice are viable, fertile and born in Mendelian ratio (supplemental Table 1). There were no obvious signs of blood lymphatic defects and both platelet and megakaryocyte counts were comparable to wildtype (WT) mice (supplemental Table 2 and supplemental Figure 2). This is in contrast to other mouse lines in which CLEC-2 or podoplanin have been genetically modified^**3,4,12**^ and suggests that human CLEC-2 can compensate for loss of the mouse protein and the interaction with murine podoplanin is sufficient for blood-lymphatic vessel separation. Human but not mouse CLEC-2 could be detected on platelets from hCLEC-2^KI^ mice with heterozygotes expressing half each of human and mouse CLEC-2 (Figure 1A and supplemental Figure 3). Surface abundance of CLEC-2 on hCLEC-2^KI^ platelets was approximately double that on human platelets (Figure 1A). The surface abundance of all other glycoprotein receptors was comparable to WT platelets (supplemental Table 3) as were platelet activation and aggregation for G protein-coupled receptors (GPCR) as well as GPVI agonists (Figure 1B-D, supplemental Figure 4). However, there was a slight increase in lag time prior to aggregation with rhodocytin, similar to that seen in human platelets (Figure 1B). Thrombus formation on collagen under flow at 1200 s^-1^ was unaltered in hCLEC-2^KI^ mice (supplemental Figure 5). Platelet spreading was comparable to WT on both fibrinogen and CRP. However, it was slightly reduced on mouse podoplanin for hCLEC-2^KI^ platelets as, although they formed both filopodia and lamellipodia, there were few fully spread platelets (supplemental Figure 6). Overall, this suggests that there are no major differences in platelet function in hCLEC-2^KI^ compared to WT mice.

**Figure 1.**
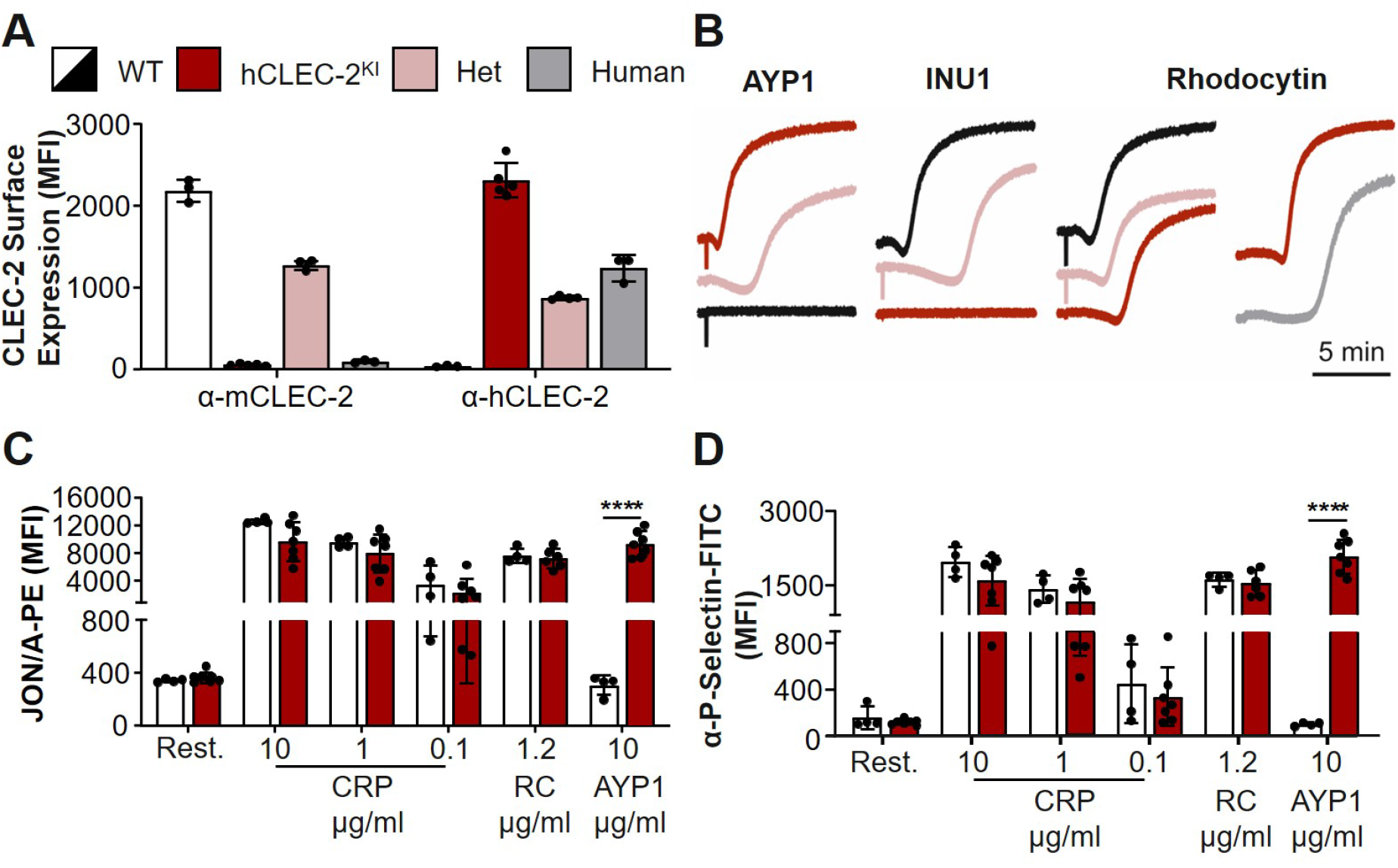
hCLEC-2^KI^ >mice have normal CLEC-2 expression and platelet activation. (**A**) Surface expression of mouse and human CLEC-2 by flow cytometry using INU1 and AYP1 antibodies respectively. Heterozygous mice expressed half human and half mouse CLEC-2 whereas hCLEC-2^KI^ mice expressed only hCLEC-2 on their platelet surface. (**B**) Light transmission aggregometry with washed platelets shows that AYP1 (10 µg/ml) and INU1 (10 µg/ml) cause aggregation of hCLEC-2^KI^ and WT platelets respectively and reduced aggregation in hCLEC-2^KI^ heterozygous platelets. Rhodocytin (0.24 µg/ml) induced aggregation has a longer lag time in hCLEC-2^KI^ platelets but also in human platelets. (**C**) Platelet integrin activation measured by JON/A-PE antibody binding in flow cytometry shows no difference in (hem)ITAM-mediated platelet activation in hCLEC-2^KI^ platelets and hCLEC-2 specific activation by AYP1. (**D**) Platelet granule secretion measured using an anti-P-selectin antibody in flow cytometry was unaltered in hCLEC-2^KI^ following platelet activation by (hem)ITAM agonists whereas AYP1 causes hCLEC-2 specific granule secretion. Data analysed by two-way ANOVA followed by a Sidak’s multiple comparison test. ****, *P* < 0.001. Results are shown as mean ± standard deviation with each circle representing one individual and are representative of three independent experiments. Het, heterozygous; MFI, mean fluorescent intensity; RC, rhodocytin; CRP, collagen related peptide; PE, phycoerythrin; FITC, fluorescein isothiocyanate.

The novel antibody HEL1 is specific to hCLEC-2 and can be used in flow cytometry, western blotting and for immunoprecipitation (supplemental Figure 7A-C). It binds to a different epitope on CLEC-2 than AYP1 as no competition between the two antibodies was observed (supplemental Figure 7D), although both antibodies cause hCLEC-2^KI^ platelet aggregation (Figure 1B and supplemental Figure 7E). Furthermore, HEL1 Fab fragments neither block rhodocytin induced platelet aggregation, unlike AYP1 Fab fragments,^14^ nor AYP1 IgG induced aggregation of hCLEC-2^KI^ platelets (supplemental Figure 7F). This not only suggests AYP1 and HEL1 act at different sites on CLEC-2 but also that CLEC-2 dimerisation, independent of its active site, is sufficient to trigger platelet activation. To investigate whether human, like mouse, CLEC-2 can be immunodepleted *in vivo*, AYP1 or HEL1 were injected intraperitoneally and CLEC-2 surface expression was determined by flow cytometry. Both antibodies deplete CLEC-2 for at least 11 days with levels returning to normal by 18 days for AYP1 and 24 days for HEL1 (Figure 2B and supplemental Figure 8). In both cases there was a transient thrombocytopenia lasting up to 4 days (Figure 2A). This is consistent with immunodepletion of mCLEC-2 using INU1 however, the length of the CLEC-2 depletion is prolonged (Figure 2C).^8,18^

**Figure 2.**
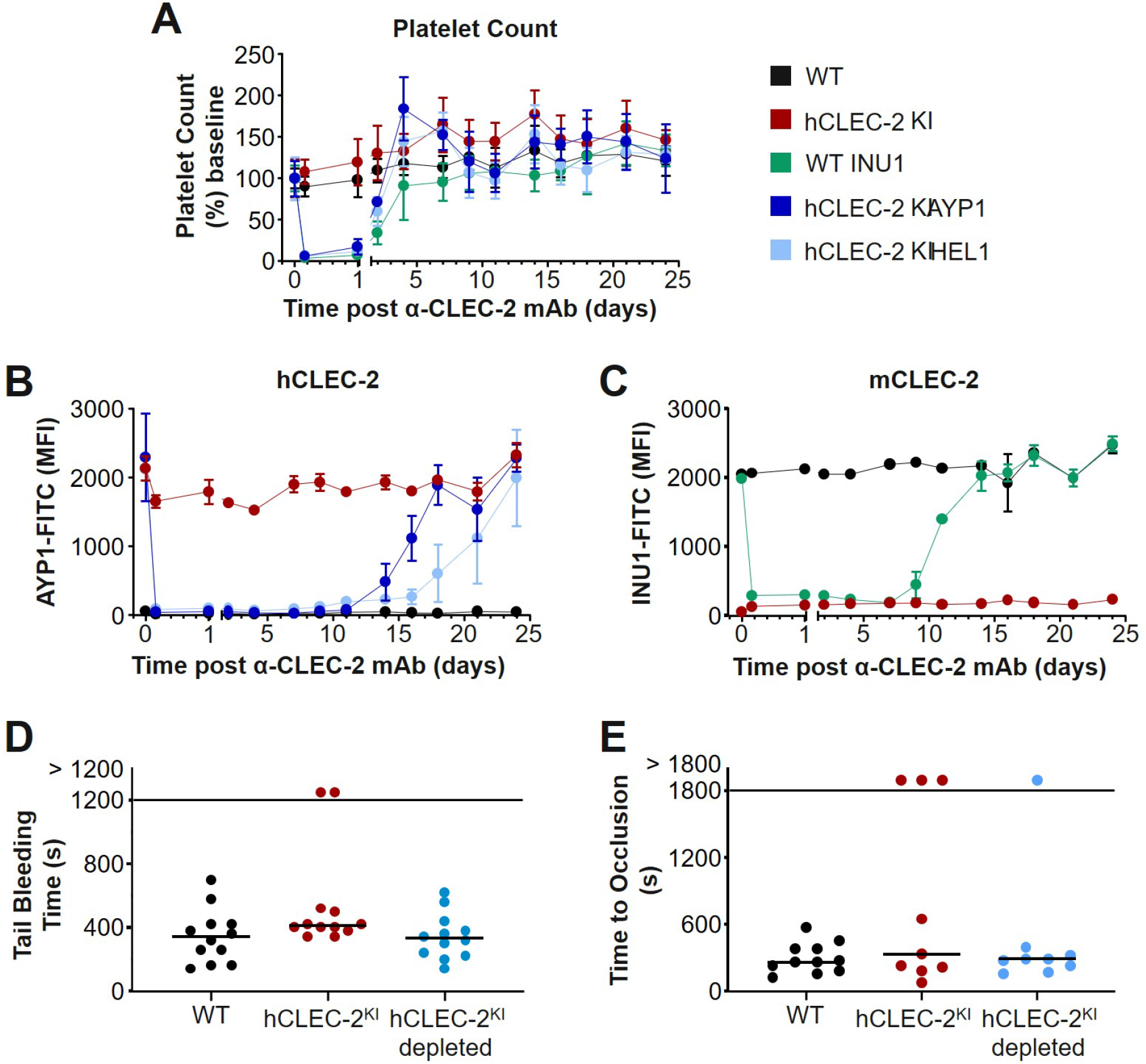
hCLEC-2 can be immunodepleted using HEL1 or AYP1 with little effect on haemostasis. **(A)** Platelet count following intraperitoneal injection of either INU1, AYP1 or HEL1 antibody (3 µg/g bodyweight). Transient thrombocytopenia lasting up to 4 days after injection can be seen for all antibody treated groups compared to the untreated controls. Platelet count was determined by flow cytometry and is shown as the percentage of the baseline count. (**B**) hCLEC-2 surface expression determined by flow cytometry following depletion by either AYP1 or HEL1. For both antibodies, CLEC-2 could not be detected on the platelet surface for at least 11 days. (**C**) mCLEC-2 surface expression determined by flow cytometry following depletion by INU1 shows CLEC-2 was absent for at least 7 days after injection. Measurements with anti-rat and anti-mouse IgG excluded the possibility that the abolished anti-CLEC-2-FITC binding was due to the presence of remaining anti-CLEC-2 antibodies on the platelets (supplemental Figure 8). (**D**) Depletion of CLEC-2 had no effect on tail bleeding time. P > 0.05, Fisher exact test. Horizontal lines represent the median time to cessation of bleeding with each circle representing one mouse. (**E**) Depletion of CLEC-2 had no effect on occlusive thrombus formation following mechanical injury of the abdominal aorta and hCLEC-2^KI^ mice were comparable to WT. P > 0.05, Fisher exact test. Horizontal lines represent the median time to vessel occlusion with each circle representing one mouse. For all experiments a minimum of 5 mice were tested per group. mAb, monoclonal antibody; FITC, fluorescein isothiocyanate; MFI, mean fluorescent intensity.

The effect of hCLEC-2 depletion was investigated in *in vivo* thrombosis and haemostasis models. Depletion had no effect on tail bleeding time (Figure 2D) which adds further support for CLEC-2 having a minor role in haemostasis.^11^ In the mechanical injury of the abdominal aorta thrombosis model there was no difference in the time to vessel occlusion in CLEC-2 depleted hCLEC-2^KI^ mice compared to untreated controls and neither group differed significantly from WT mice (Figure 2E). Similar results have been shown in platelet specific CLEC-2 knockout mice^12^. However, a reduction in occlusive thrombus formation has been observed in other models of thrombosis suggesting the contribution of CLEC-2 to thrombosis differs depending on the type of injury or the vascular bed.^8,11,12^ Notably, hCLEC-2 can compensate for mCLEC-2 during development as well as in thrombosis and haemostasis suggesting a conserved interaction between CLEC-2 and podoplanin.

Here, we have demonstrated that hCLEC-2 can be immunodepleted and provide proof of principle that the hCLEC-2^KI^ mouse can be used to test anti-hCLEC-2 agents *in vivo*.

## Acknowledgements

This project has received funding from the European Union’s Horizon 2020 research and innovation programme under the Marie Skłodowska-Curie grant agreement No 766118. This work was supported by TR240 grant with project number 374031971 of the Deutsche Forschungsgemeinschaft (DFG; German Research Foundation) and the Rudolf Virchow Center. We thank Stefanie Hartmann, Juliana Goldmann and Ewa Stepien-Bötsch for excellent technical assistance.

## Authorship

Contributions: HCB performed experiments, analysed data and wrote the manuscript; SB, SN, EMSJ and JK preformed experiments and analysed data; YD and VM provided vital reagents and proofread the manuscript; ST, SPW and BN supervised the research and wrote the manuscript; DS supervised the research, performed experiments, analysed the data and wrote the manuscript.

## Supplemental Tables

**Supplemental Table 1.** Analysis of genotypes at weaning of pups from heterozygous hCLEC-2^KI^ matings.

|  | Expected | Observed |
| --- | --- | --- |
| **Wildtype** | 52 | 63 |
| **Heterozygous** | 102 | 92 |
| **hCLEC-2<sup>Kl</sup>** | 52 | 51 |
| **Total** | 206 | 206 |
| **Chi<sup>2</sup>** |  | 1.58 (P = 0.45) |

**Supplemental Table 2.**
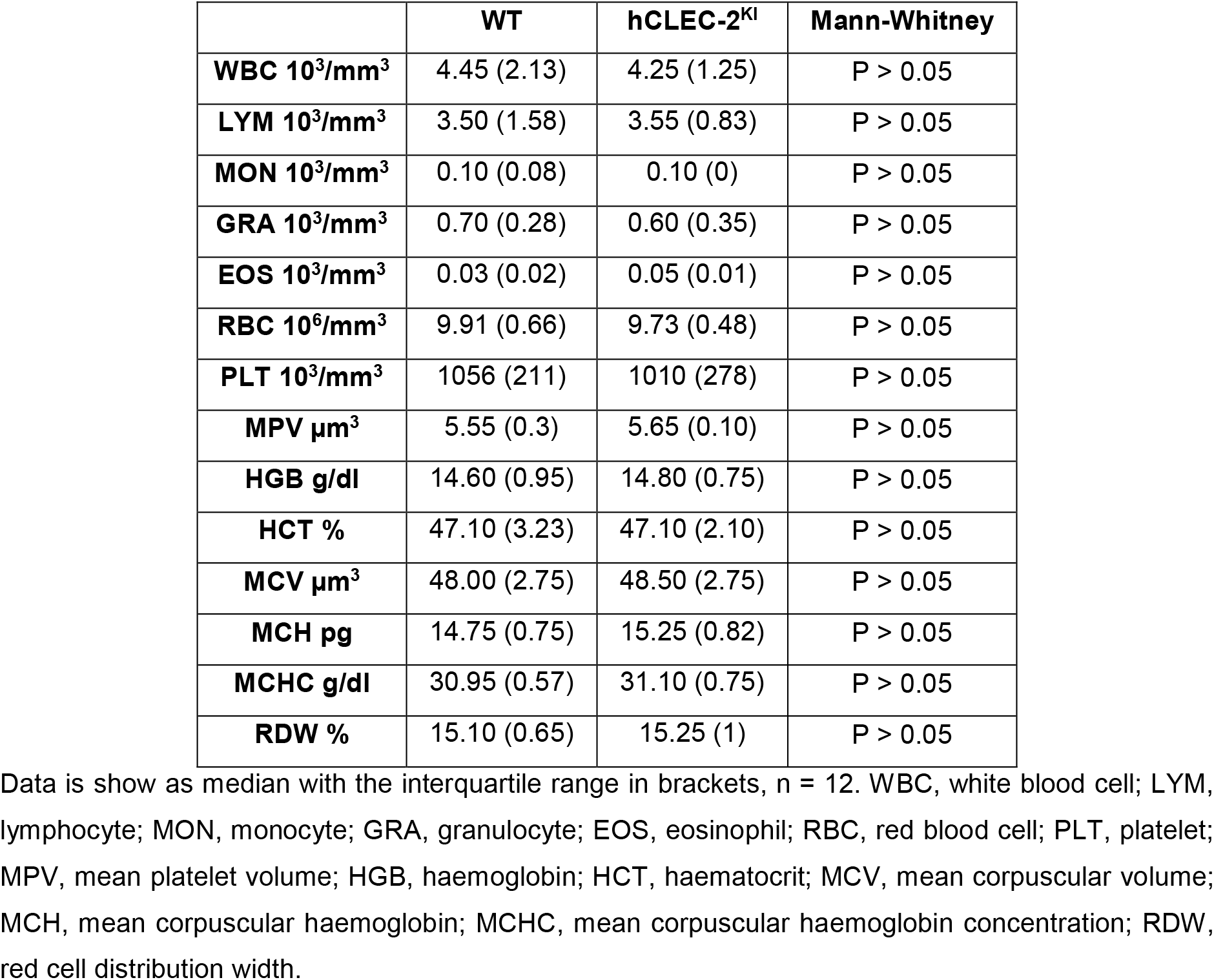
Analysis of blood parameters shows no difference between wildtype and hCLEC-2^KI^ mice.

**Supplemental Table 3.**
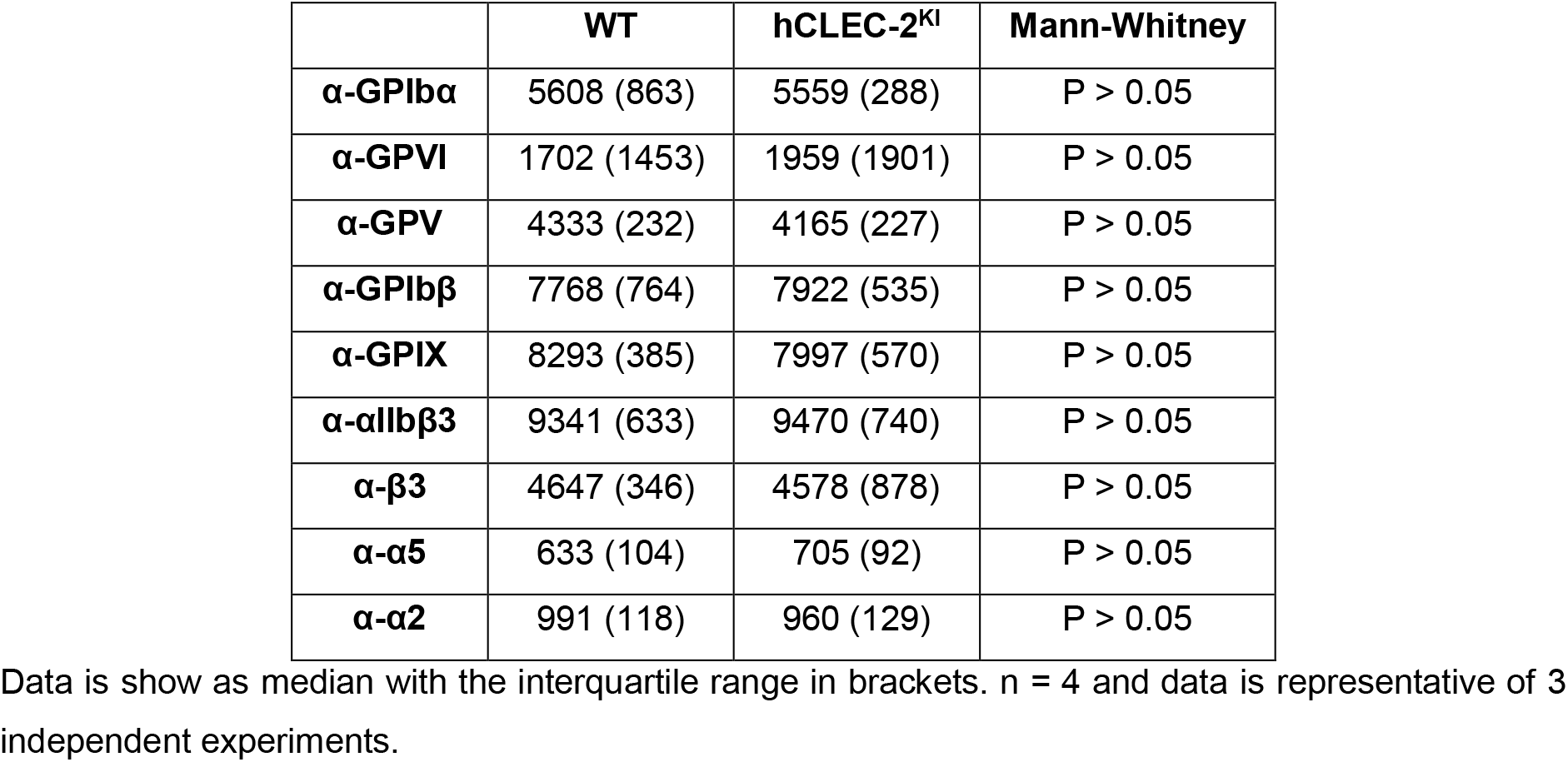
Comparison of platelet glycoprotein expression shows no difference between wildtype and hCLEC-2^KI^ mice.

## Supplemental Figures

**Supplemental Figure 1.**
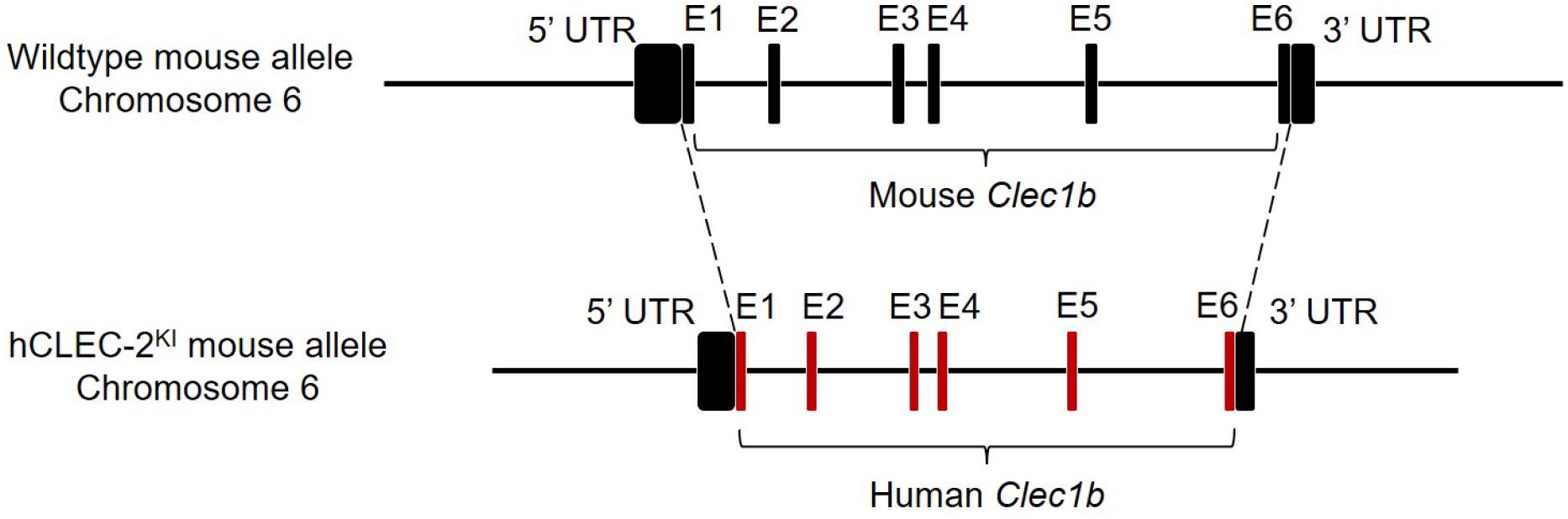
Generation of hCLEC-2^KI^ mice. CRISPR/Cas9 was used to replace the wildtype mouse *Clec1b* gene on chromosome 6 with the human variant. The entire mouse gene was replaced with the human but the three and five prime untranslated regions (3’ and 5’ UTR) remained as the original mouse sequence.

**Supplemental Figure 2.**
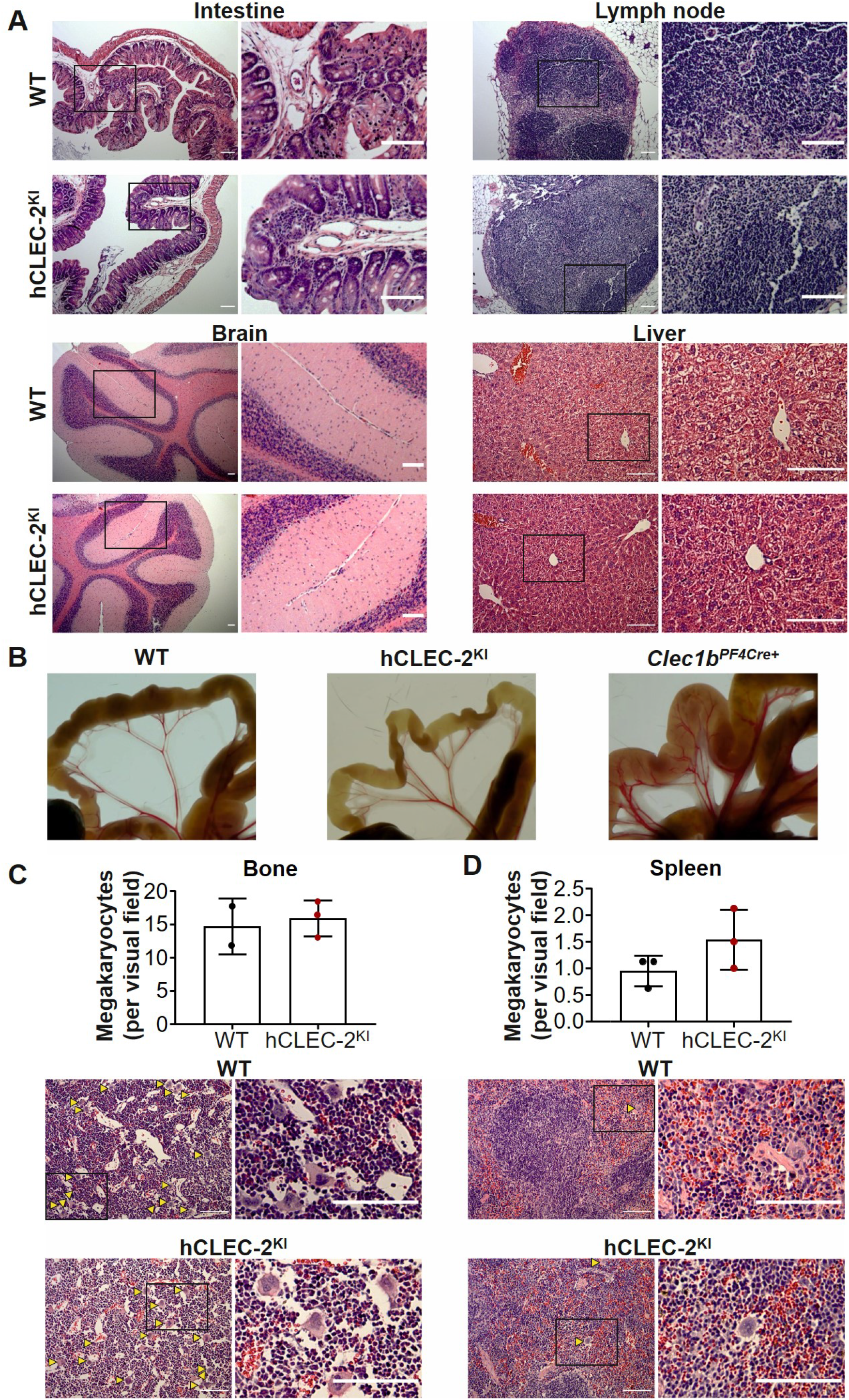
(see next page). No evidence of blood-lymphatic mixing and comparable organ morphology between hCLEC-2^KI^ and WT mice. (**A**) Haematoxylin and eosin staining of intestines, lymph nodes, brains and livers from hCLEC-2^KI^ and WT mice show no differences in morphology. (**B**) Intestines from adult WT, hCLEC-2^KI^ and *Clec1b*^*PF4Cre+*^ mice were compared for evidence of blood-lymphatic mixing. In *Clec1b*^*PF4Cre+*^ but not hCLEC-2^KI^ mice there is increased blood in the mesenteric vessels. (**C**) Quantification of megakaryocytes in haematoxylin and eosin stained bone sections shows no difference between hCLEC-2^KI^ and WT mice or in bone morphology. (**D**) Quantification of megakaryocytes in haematoxylin and eosin stained spleen sections show no difference between hCLEC-2^KI^ and WT mice or in spleen morphology. Scale bars represent 100 µM. Yellow arrowheads indicate megakaryocytes. Images are representative of n = 3. Results are shown as mean ± standard deviation.

**Supplemental Figure 3.**
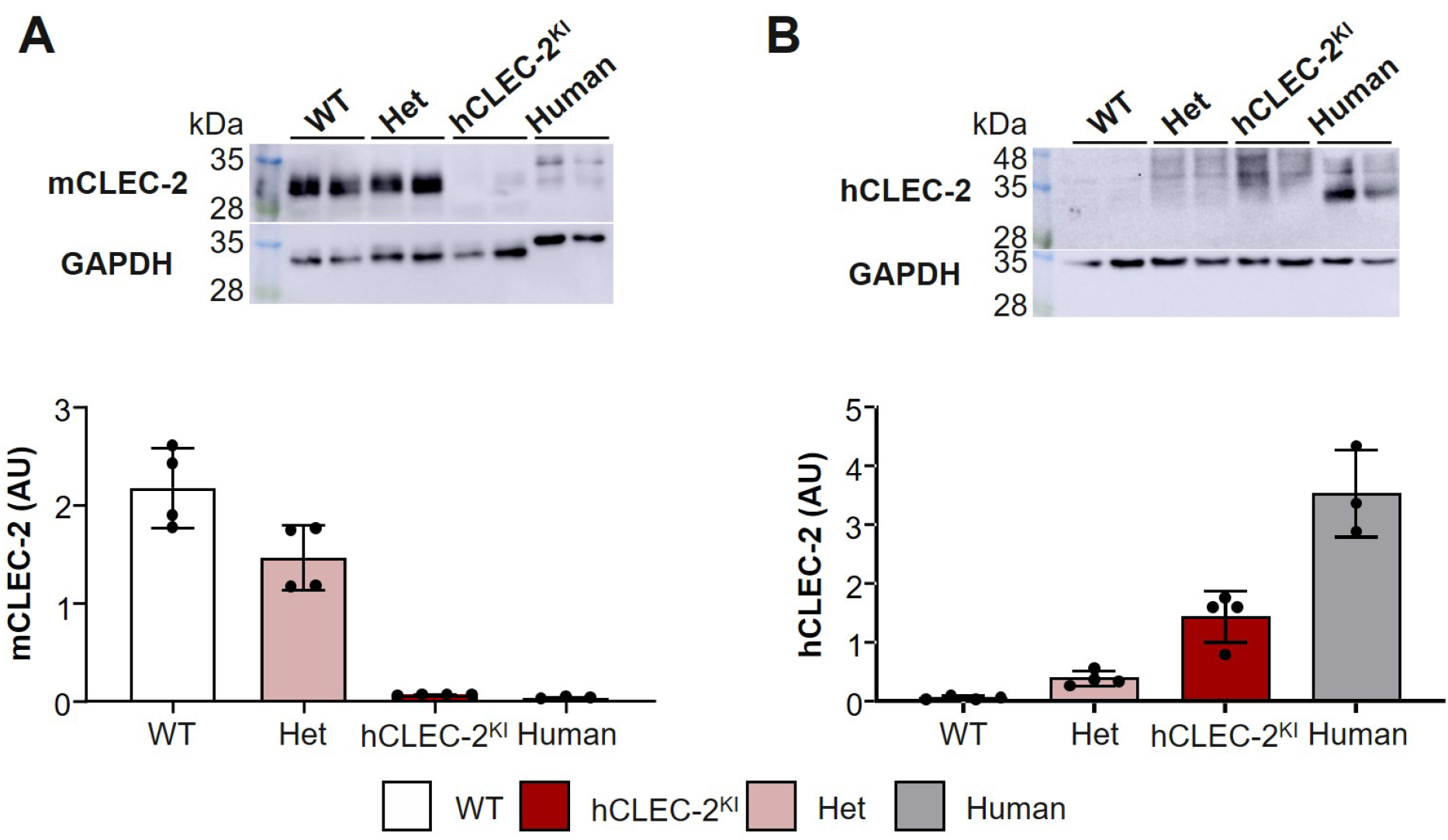
hCLEC-2^KI^ mice only express human CLEC-2. Western blot analysis of platelet lysates from WT, heterozygous, hCLEC-2^KI^ and healthy human donors show hCLEC-2^KI^ mouse platelet express human but not mouse CLEC-2. CLEC-2 expression was normalised to the loading control GAPDH. (**A**) Representative Western blot and quantification of mouse CLEC-2 using the antibody INU2. (**B**) Representative Western blot and quantification of human CLEC-2 using the antibody AYP2. Results are shown as mean ± standard deviation and each circle represents one individual. Het, heterozygous; GAPDH, Glyceraldehyde 3-phosphate dehydrogenase; AU, arbitrary units.

**Supplemental Figure 4.**
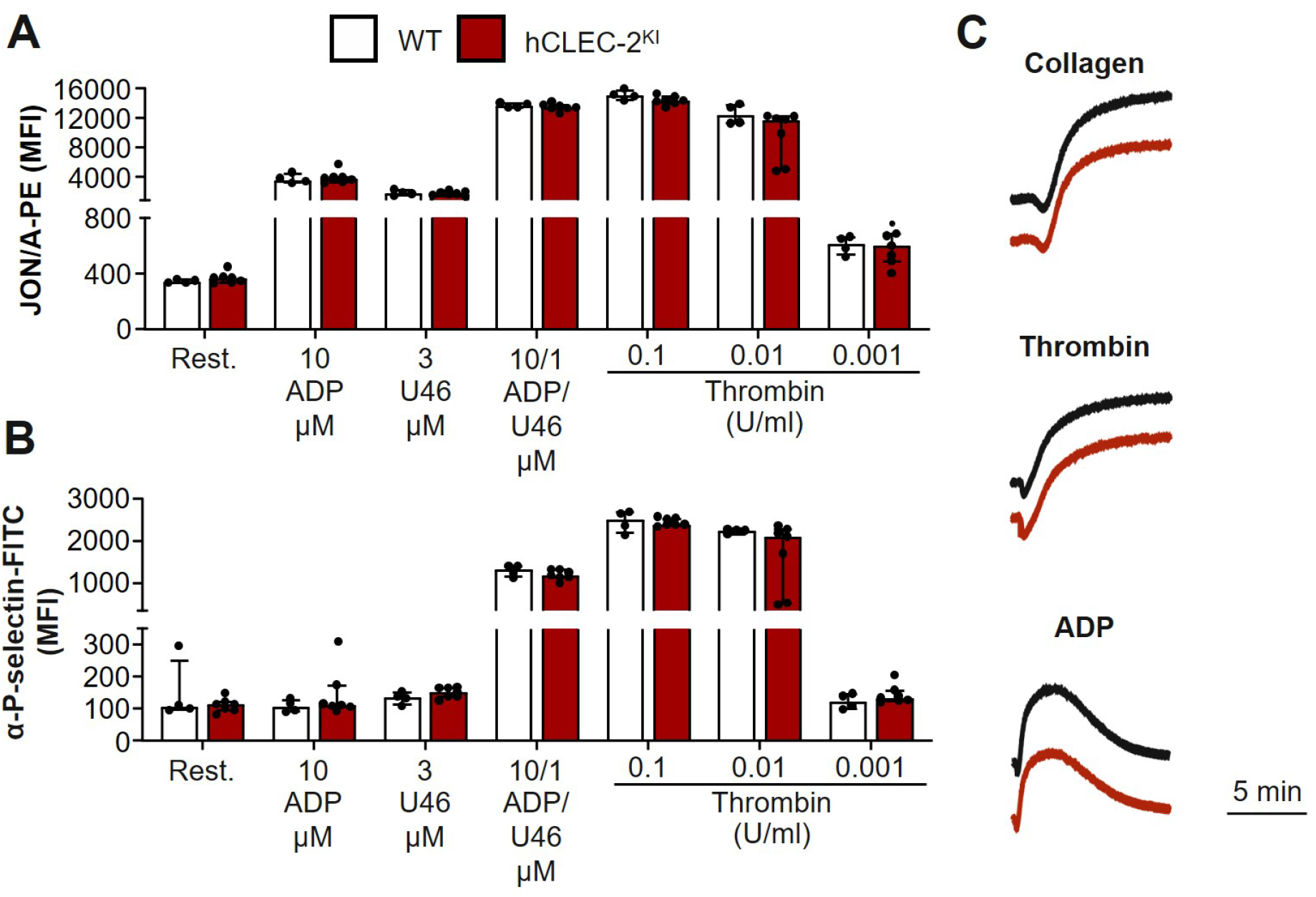
hCLEC-2^KI^ mice have normal platelet activation and aggregation. (**A**) Platelet integrin activation measured by JON/A-PE binding in flow cytometry shows no difference in hCLEC-2^KI^ compared to WT mice following stimulation with ADP, U46619 or thrombin. Data were analysed using Mann-Whitney tests (P > 0.05) (**B**) Platelet granule secretion measured using an anti-P-selectin antibody in flow cytometry was unaltered in hCLEC-2^KI^ following platelet activation by ADP, U46619 or thrombin. Data were analysed using Mann-Whitney tests (P > 0.05) (A, B) each circle represents one individual. Results are shown as mean ± standard deviation. (**C**) Light transmission aggregometry with washed platelets shows unaltered aggregation in hCLEC-2^KI^ mice in response to collagen (5 µg/ml) or thrombin (0.01 U/ml) as well as no difference in ADP (5 µM) induced aggregation in platelet rich plasma. Traces are representative of at least n = 3. PE, phycoerythrin; MFI, mean fluorescent intensity; Rest, resting; ADP, adenosine diphosphate; U46, thromboxane A2 mimetic U46619; FITC, fluorescein isothiocyanate.

**Supplemental Figure 5.**
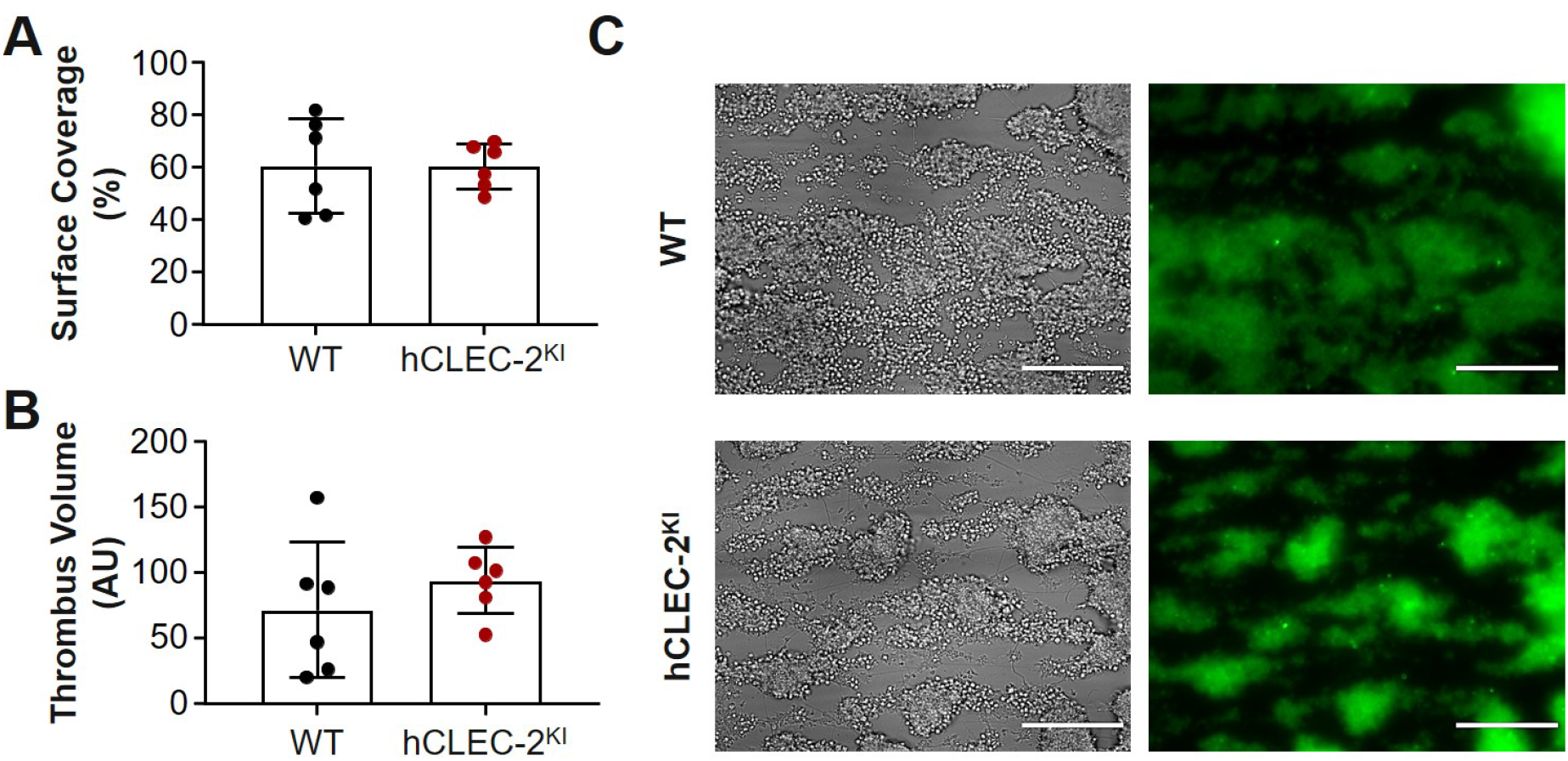
Thrombus formation on collagen under flow is unaltered in hCLEC-2^KI^ mice at a shear rate of 1200 s^-1^. (**A**) Surface coverage of thrombi formed on collagen was comparable between WT and hCLEC-2^KI^ mice (unpaired t-test, P = 0.99). (**B**) Thrombus volume did not differ between WT and hCLEC-2^KI^ mice (unpaired t-test, P = 0.36). (**C**) Representative brightfield and fluorescent images of aggregates formed under flow on collagen from WT and hCLEC-2^KI^ mice. Platelets were labelled prior to flow with an anti-GPIX-Dylight488 conjugated antibody. Scale bars represent 50 µm. Data are shown as mean ± standard deviation and each circle represents one animal. AU, arbitrary units.

**Supplemental Figure 6.**
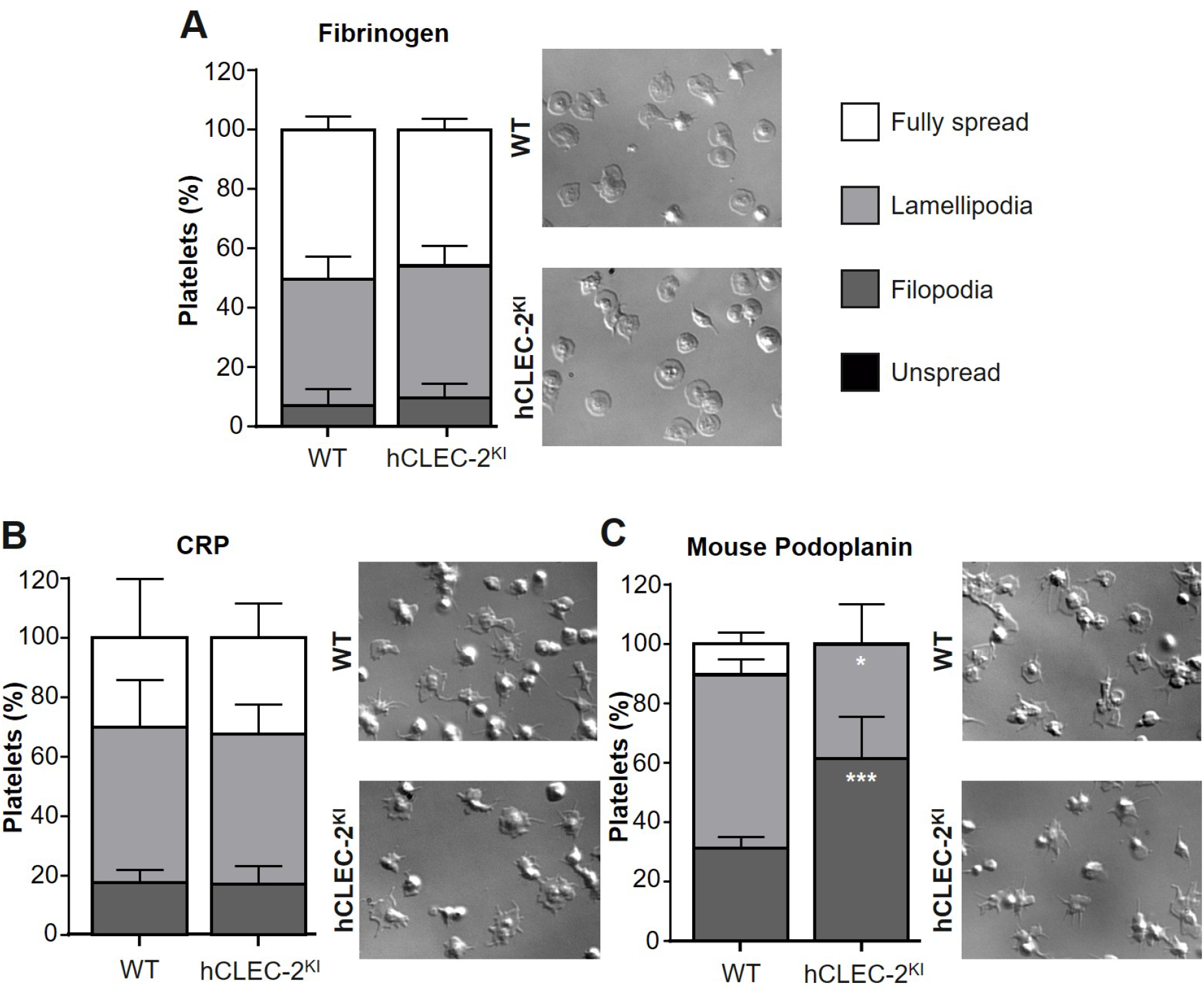
Platelet spreading is unaltered on fibrinogen and CRP but reduced on mouse podoplanin in hCLEC-2^KI^ mice. Platelet spreading is shown as the percentage of platelets in each of the 4 stages of spreading: unspread, forming filopodia, forming lamellipodia and fully spread platelets from 5 fields of view plus representative images. (**A**) Platelet spreading on 100 µg/ml human fibrinogen was comparable between WT and hCLEC-2^KI^ mice after 30 min (P = 0.63). 0.01 U/ml thrombin were added to the platelets immediately prior to spreading. (**B**) Platelet spreading on 10 µg/ml CRP after 30 min was comparable for WT and hCLEC-2^KI^ mice (P = 0.99). (**C**) Platelet spreading on 10 µg/ml recombinant mouse podoplanin-FC was reduced in hCLEC-2^KI^ mice compared to WT. More platelets had filopodia (P = 0.0006) in the hCLEC-2^KI^ group and fewer formed lamellipodia (P = 0.0188). However, no unspread platelets were observed. Data was analysed using a two-way ANOVA with Sidak’s multiple comparison test. Data represents 3 mice per genotype and is representative of at least 2 independent experiments. *** P < 0.001, * P < 0.05.

**Supplemental Figure 7.**
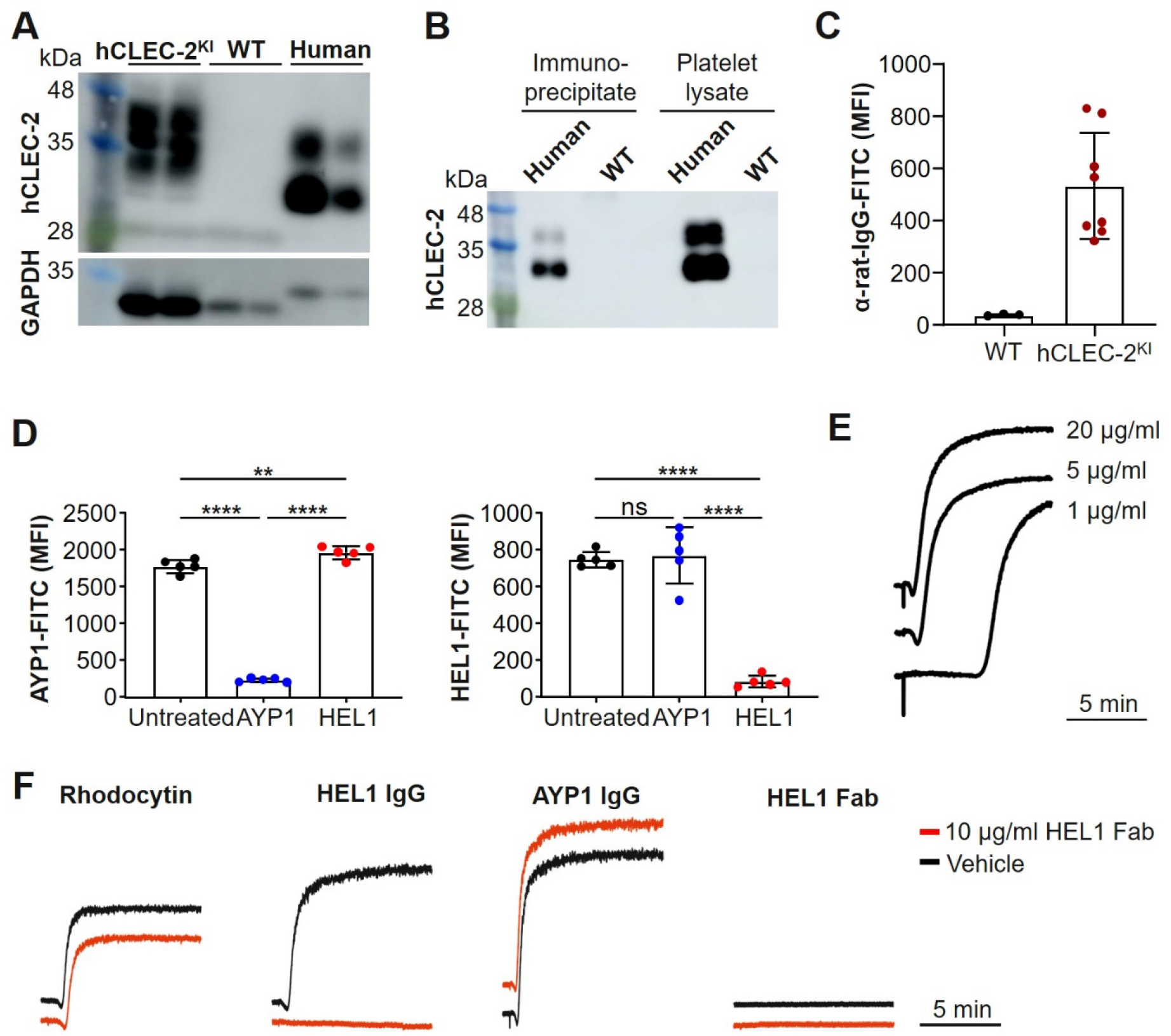
HEL1 is specific to hCLEC-2 and appears to bind at a different site to AYP1 but still activates hCLEC-2^KI^ platelets. (**A**) HEL1 can detect hCLEC-2 from both hCLEC-2^KI^ and human platelet lysates but not WT mouse lysates. (**B**) HEL1 immunoprecipitates hCLEC-2. This is shown by Western blotting of the eluate following immunoprecipitation by HEL1 (5 µg) coupled to protein G Sepharose beads from platelet lysate. (**C**) HEL1 can detect hCLEC-2 on the surface of hCLEC-2^KI^ but not WT platelets by flow cytometry. HEL was incubated with whole blood, which was then washed by centrifugation and an anti-rat IgG-FITC coupled antibody was used to detect HEL1 on the platelet surface. (**D**) HEL1 and AYP1 bind to different sites on hCLEC-2 as shown by flow cytometry. hCLEC-2^KI^ mouse blood incubated with 10 µg/ml AYP1 had normal surface expression of hCLEC-2 determined using HEL1-FITC and blood incubated with 10 µg/ml HEL1 had normal hCLEC-2 surface expression when measured with AYP1-FITC. In both cases incubating and measuring hCLEC-2 surface expression with the same antibody resulted in a reduction in expression compared to an untreated control due to opsonization of the respective binding site. (**E**) HEL1 IgG causes dose-dependent aggregation of hCLEC-2^KI^ platelets. (**F**) HEL1 Fab fragments do not block rhodocytin (0.24 µg/ml) or AYP1 IgG (10 µg/ml) induced hCLEC-2^KI^ platelet aggregation but do prevent HEL1 IgG (10 µg/ml) induced aggregation providing further evidence HEL1 and AYP1 bind to different epitopes on hCLEC-2. HEL1 Fab fragments themselves do not induce platelet aggregation. Data are show as mean ± standard deviation and statistical significance was determined by one-way ANOVA followed by Tukey’s multiple comparisons test. ****, P < 0.001 **, P < 0.01. Each circle represents one individual. MFI, mean fluorescent intensity; FITC, fluorescein isothiocyanate.

**Supplemental Figure 8.**
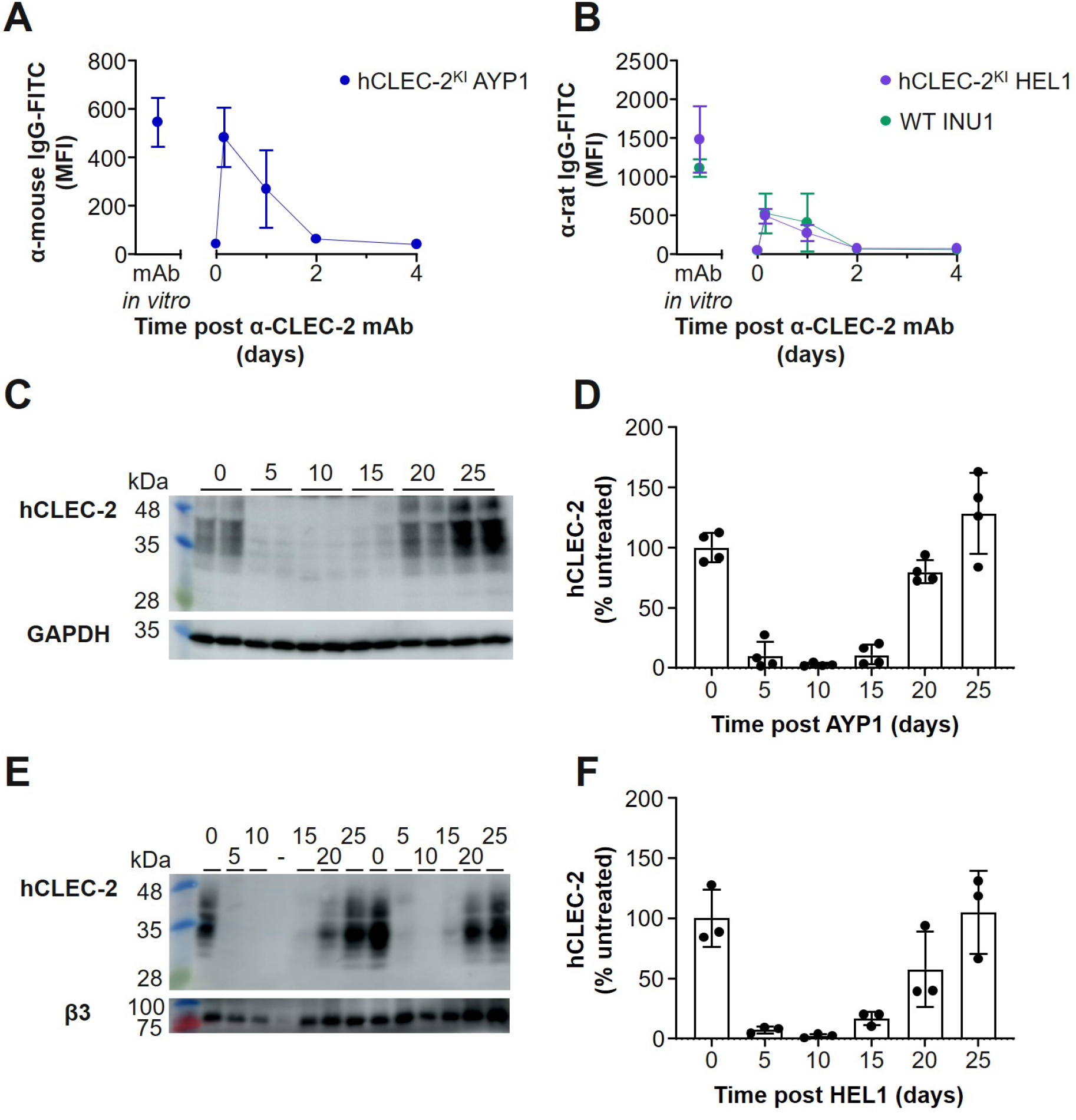
hCLEC-2 immunodepletion by HEL1 and AYP1 results in CLEC-2 deficient platelets. (**A**) AYP1 can be detected on the surface of platelets after intraperitoneal injection using an anti-mouse IgG-FITC conjugated antibody for only 1 day following injection. This is shown as an increased in MFI following antibody injection compared to the baseline measurements. (**B**) Traces of HEL1 and INU1 can be detected on the surface of hCLEC-2^KI^ and WT platelets respectively for only 1 day after intraperitoneal injection using an anti-rat IgG-FITC conjugated antibody. This is shown as an increased in MFI following antibody injection compared to the baseline measurements. (**C**) Intraperitoneal AYP1 (3 µg/g bodyweight) depletes both intra- and extracellular hCLEC-2. Western blotting of platelet lysates from AYP1 injected mice at the various time points using AYP2 shows that hCLEC-2 is absent from platelets between 5 and 15 days after injection, earlier time points could not be studied due to the thrombocytopenia caused by AYP1 injection. (**D**) Quantification of Western blots showing hCLEC-2 depletion following AYP1 injection. hCLEC-2 levels increase slightly at 15 days after injection and return fully between day 20 and 25. Data is shown as the percentage of hCLEC-2 compared to untreated mice and normalised to the loading control GAPDH. (**E**) Intraperitoneal HEL1 (3 µg/g bodyweight) depletes both intra- and extracellular hCLEC-2. Western blotting of platelet lysates from HEL1 injected mice at the various time points using HEL1 shows that hCLEC-2 is absent from platelets between 5 and 15 days after injection, earlier time points could not be studied due to the thrombocytopenia caused by HEL1. “-” indicates an empty lane. (**F**) Quantification of Western blots showing hCLEC-2 depletion following HEL1 injection. hCLEC-2 levels increase slightly 15 days after injection and return fully between day 20 and 25. Data is shown as the percentage of hCLEC-2 compared to untreated mice and normalised to the loading control integrin β3. MFI, mean fluorescent intensity; GAPDH, Glyceraldehyde 3-phosphate dehydrogenase.

